# Ovarian Hormones Moderate Systolic Hypertension in Female *Eln* Haploinsufficient Mice

**DOI:** 10.1101/2025.11.18.689164

**Authors:** Alethia J. Dixon, Ipsita Mohanty, Gagandeep Kaur, James A. McCormick, Patrick Osei-Owusu

**Affiliations:** Department of Physiology & Biophysics, Case Western Reserve University School of Medicine, Cleveland, Ohio, United States; Department of Pharmacology and Physiology, Drexel University College of Medicine, Philadelphia, Pennsylvania, United States; Department of Medicine, Oregon Health & Science University, Portland, Oregon, United States

**Keywords:** elastin haploinsufficiency, hypertension, ovarian hormones, renal function, vasopressin signaling

## Abstract

Hypertension is a hallmark of cardiovascular abnormalities associated with Williams syndrome (WS), a rare genetic disorder involving microdeletion of genes on human chromosome 7, including the elastin gene (*ELN*). Heterozygous deletion of *Eln* (*Eln^+/−^*) in mice recapitulates hypertension and arteriopathy associated with WS. Previously, differences in blood pressure elevation and sensitivity to dietary sodium were found to be less profound in female *Eln^+/−^* mice. Here, we determined whether ovarian hormones play a role in sex-related difference in blood pressure elevation resulting from *Eln* haploinsufficiency. Female *Eln^+/+^* and *Eln^+/−^* mice instrumented with radiotelemetry devices were subjected to sham surgery or ovariectomy (OVX). We found that OVX lowered diastolic but not systolic blood pressure (SBP) in *Eln^+/−^* mice, resulting in increased pulse pressure. In *Eln^+/−^* mice, diuresis induced by acute volume expansion was blunted, while anti-natriuresis was exaggerated. Furthermore, amiloride lowered SBP and increased urinary Na^+^ excretion, suggesting that *Eln^+/−^*-induced hypertension may be Na^+^-dependent. We conclude that increased Na^+^ and water retention by the kidney contribute to hypertension resulting from *Eln* haploinsufficiency. The underlying mechanism involves the alteration of ovarian hormone effects in the kidney and sustained signaling downstream of the V_2_ receptor, leading to increased ENaC activity and water reabsorption.

Graphical Abstract
CDPC, cortical duct principal cell; ER, estrogen receptor; PR, progesterone receptor; ENaC, epithelial sodium channel; AQP2, aquaporin 2; V_2_R, vasopressin V_2_ receptor; ELN, elastin; PKA, protein kinase A; AC, adenylyl cyclase; Gs, stimulatory G protein; ATP, Adenosine triphosphate; cAMP, cyclic AMP; ⊕, activating; ⊖, inhibiting

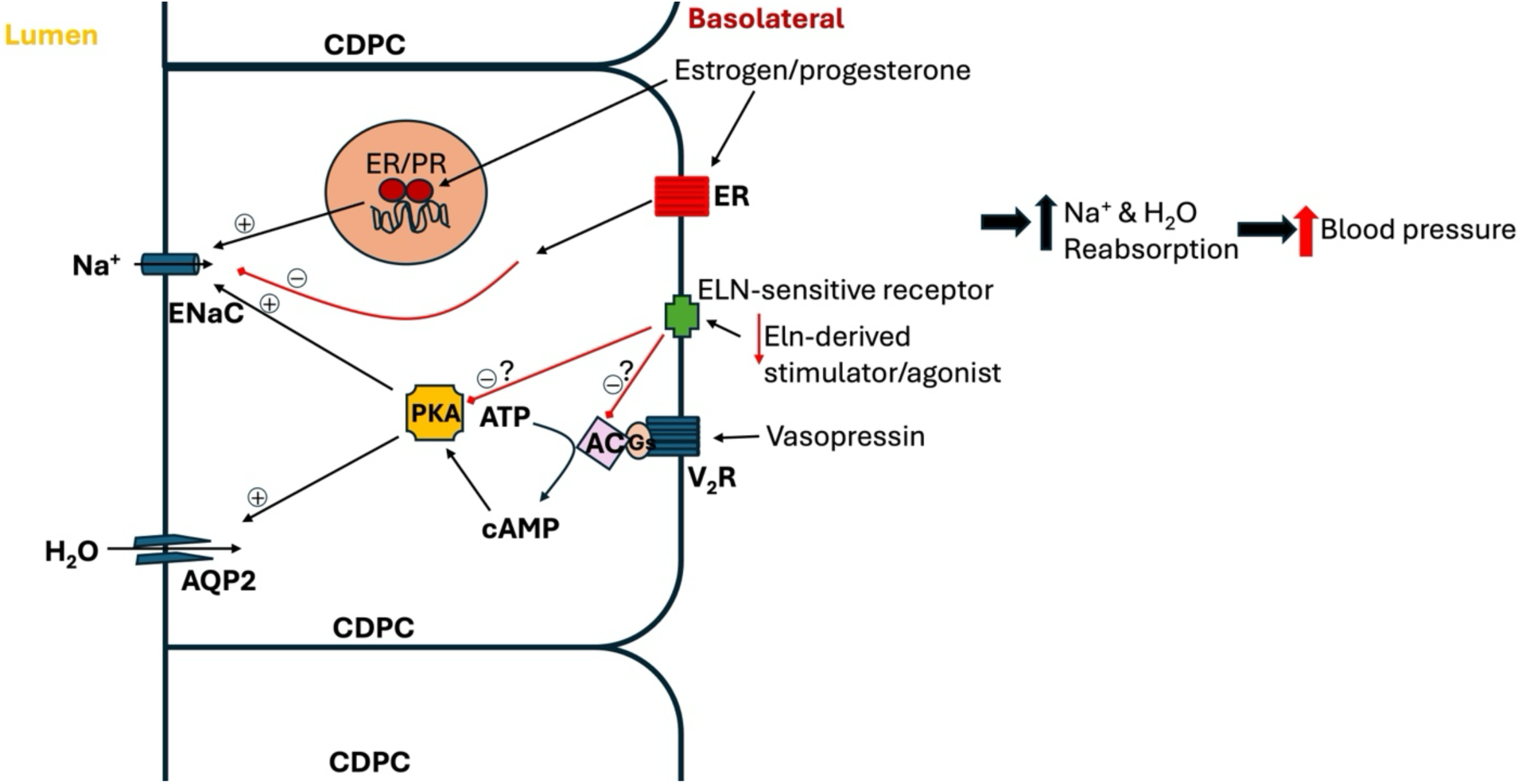

## Introduction

Hypertension is a leading risk factor for the development of cardiovascular disorders and chronic kidney disease and is notably more prevalent in post-menopausal women but less in pre-menopausal women compared to age-matched males^1,2^. Several studies establish the anti-hypertensive effects of ovarian hormones as a primary biological difference between males and females^3,4^. Ovarian hormones, particularly estrogen, modulate cardiovascular function by activating endothelial nitric oxide synthase (eNOS) and influencing the activity of the renin-angiotensin system (RAS)^5–8^. Furthermore, estrogen receptors localized on renal epithelial cells regulate the activity and expression of sodium channels and transporters along the nephron^9–11^, suggesting a role for sex hormones in the regulation of fluid and electrolyte balance, which serves as a primary mechanism by which the kidney maintains long-term blood pressure control. However, the precise mechanisms by which sex hormones confer such protective effects, either by acting in the kidney or in other organ systems, are incompletely understood.

Long-term blood pressure control by the kidney is a complex physiological process involving the regulation of renal hemodynamics and controlled tubular reabsorption to maintain extracellular fluid volume and electrolyte balance^12,13^. Central to this process is renal autoregulation, which is mediated by kidney intrinsic mechanisms that are in turn finetuned by factors that are either intrinsic or extrinsic to the kidney. Renal autoregulation is key to adjusting renal vascular resistance (RVR) to stably maintain renal blood flow (RBF) and glomerular filtration rate (GFR), despite fluctuating changes in systemic blood pressure^14,15^. Abnormal autoregulation can potentially lead to renal ischemia or barotraumatic damage of the renal microvasculature and the glomerular capillary network^14,16^. Such damage and autoregulation failure could lead to hyperfiltration and over-saturation of the renal tubular system, thereby causing proteinuria and/or inappropriate loss or excessive retention of electrolytes and water sustaining hypertension^15,17^.

Chronic hypertension elevates renal perfusion pressure and is associated with elevated peripheral vascular resistance and stiffening, that together are implicated in the impairment of renal autoregulation. Increased vascular stiffness develops partly from the loss of functional elastin (ELN), the principal extracellular matrix (ECM) protein in vascular elastic fibers that form the internal, medial, and external elastic laminae^18,19^. ELN is produced and secreted into the ECM as tropoelastin monomers, which concatenate in structured fibrillar networks of other ECM proteins and are finally crosslinked by lysyl oxidase to form mature elastin fibers^20,21^. Mature ELN is formed in early life but fatigues with age and is also gradually fragmented and degraded by secreted elastases and local matrix metalloproteinases (MMPs), thus progressively losing its initial mechanical properties without a replacement^22–24^. ELN degradation also increases collagen (COL) deposition, thereby increasing COL/ELN ratio^22,23,25^. As such, disease– and age-related vascular remodeling, including arterial stiffening, calcification, and dysfunction are also associated with deficiency of functional ELN and tissue fibrosis^26–28^.

Previously, we reported that *Eln* haploinsufficiency (*Eln*^+/−^) in mice is independently associated with increased RVR and pathological remodeling of the renal microvsculature^29^. These structural and functional changes were shown to precede the onset of hypertension that becomes fully established in *Eln*^+/−^ mice by 1 month of age^30^. Furthermore, the hypertensive phenotype in *Eln*^+/−^mice was found to be more pronounced in male than female mice, when compared to their respective *Eln*^+/+^ controls, suggesting a potential role for sex-related differences, including differences in sex hormone levels, in mitigating or driving hypertension in *Eln* haploinsufficiency. However, the relationship between functional ELN level and the biological effects of sex hormones in the pathogenesis of hypertension has not been thoroughly explored.

Previously, we showed that accelerated structural changes in the renal microvasculature associated with *Eln* haploinsufficiency exacerbate age-related impairment of renal hemodynamics in female mice^31^. Whether *Eln* haploinsufficiency leads to the disruption of the homeostatic regulation of water and electrolyte balance by the kidney to contribute to chronic blood pressure elevation has not been fully explored. Furthermore, whether changes in sex hormone levels impinge on the effects of *Eln* haploinsufficiency to drive sustained blood pressure elevation is also unknown. In this study, we postulated that ovarian hormones mitigate blood pressure elevation in *Eln*^+/−^ female mice by fine-tuning renal tubular handling of water and sodium. To test this hypothesis, we investigated the effects of ovarian hormone depletion, via ovariectomy (OVX) in wild type (*Eln*^+/+^) and *Eln*^+/−^ female mice, on diurnal blood pressure and fluid and sodium handling by the kidney. Our results show that *Eln* haploinsufficiency has differential effects on the response of systolic and diastolic blood pressure to ovarian hormone depletion, which might be related to effects on renal tubular sodium and water handling.

## Materials and Methods

### Animals

The study was performed under a protocol (# 2020-0025) approved by the Institutional Animal Care and Use Committee of Case Western Reserve University. Experiments were performed using 3–4-month-old female wild-type (*Eln*^+/+^) and *elastin* heterozygous (*Eln*^+/−^) mice that have been backcrossed several generations into the C57 Bl/6 genetic background (Charles River, MA). The generation of *Eln*^+/−^ mice has been described previously^30,32^. Mice were provided access to food and water ad libitum in our institution’s animal resource center at 22°C with a 12-h light/dark cycle.

### Ovariectomy and sham procedures

All appropriate steps were taken to perform aseptic surgery for oophorectomy/ovariectomy (OVX) as described previously, with slight modifications (supplemental Figure 1A)^33–35^. Briefly, mice were shaved on the dorsal midline and an incision of ∼1 cm was made. The skin was detached from the adjoining muscle wall using blunt dissection, followed by a 10-mm incision through the abdominal wall. The ovary was externalized and removed by cauterization between the end of the uterine horn, the ovary and ovarian artery. The abdominal wall was closed with an absorbable suture and the skin closed with metal staples. The same procedure was repeated on the other side to remove the second ovary, and finally, animals were injected with meloxicam as part of post-operative pain management. Sham-operated animals were subjected to the same surgical procedure, except that the ovaries were exteriorized briefly but not excised. The success of the ovariectomy was confirmed by uterine atrophy, determined by harvesting and weighing the uterus post-euthanasia and normalizing to body weight. Ovariectomized mice with marginal uterine atrophy relative to sham controls were excluded from the study (supplemental Figure 1A – 1C). The mice were used for experiments at least 4 weeks post-ovariectomy or sham surgery.

### Diurnal blood pressure and heart rate monitoring using radiotelemetry

Conscious blood pressure data were acquired using radiotelemetry (HD-X10, Data Sciences International, St. Paul, MN), as described previously^29,36^. Briefly, the pressure-sensing catheter of the telemeter was implanted in the left carotid artery and advanced into the aortic arch, while the transmitter body was placed subcutaneously, under isoflurane anesthesia. After a 7-day recovery period, baseline blood pressure and heart rate were recorded for 24 hours. Subsequently, mice underwent ovariectomy, and blood pressure, heart rate, and locomotor activity were recorded at least 4 weeks after ovariectomy.

### Dietary sodium modification and diurnal blood pressure and heart rate recording

Sham-operated and OVX mice were placed on standard mouse chow (0.22% sodium; 5P76/P3000, LabDiet, Richmond, IN) or on a sodium-deficient diet (0.01% sodium; Invigo, cat. #: TD.90228) for 10 days, after which blood pressure and heart rate were recorded for 24 hours. The mice then received amiloride (1.5 mg/kg. i.p.) once daily for 3 days followed by 24-hour radiotelemetry recording and overnight urine collection using a mouse metabolic cage.

### Glomerular filtration rate measurement in conscious mice

Glomerular filtration rate (GFR) was assessed using the transdermal method, as previously described, with minor modifications^37^. Mice were initially implanted with a polyethylene-10 (PE-10) catheter into the jugular vein for fluorescein-isothiocyanate (FITC)-labeled sinistrin administration. The hair at the nape was shaved and disinfected, followed by small midline incisions made at the ventral and dorsal regions of the neck. A metal trocar was cautiously utilized to create a tunnel from the front to the back of the neck, through which the PE-10 catheter was threaded. The subcutaneous tissue was carefully dissected to expose the jugular vein, where the PE-10 catheter was inserted and secured with a 5-0 silk. The catheter was flushed with a sterile 10% heparinized saline solution to confirm proper placement and to prevent blood clot formation. The section of the catheter extending to the nape was sealed using a cauterizer and then secured with suture and tissue glue. To prepare the mouse for GFR measurement, the flank of the animal was shaved, followed by the application of hair removal cream before attaching the GFR mini device to an adhesive patch. The device was fastened with tape and allowed to record a steady baseline reading for ∼5 minutes before administering a bolus of FITC-sinistrin (35 mg/mL in saline) via the jugular vein catheter. Transcutaneous fluorescence was recorded for 2 hours, and the acquired data was used to calculate GFR using the manufacturer-provided software (MB_Lab2, Medibeacon). GFR values were normalized to body weight.

### Renal blood flow procedure

Simultaneous renal blood flow (RBF) and blood pressure were measured under isoflurane anesthesia in sham and OVX *Eln*^+/+^ and *Eln*^+/−^ mice using a well-established technique, as previously described^38^. Briefly, catheters were surgically implanted into the left carotid artery and right jugular vein under aseptic conditions. Following this procedure, a left flank incision was made to expose the left renal artery. A perivascular flow probe (0.5PSB Nanoprobe with handle, Transonic, Ithaca, NY) was carefully positioned around the left renal artery to enable real-time measurement of RBF. Baseline blood pressure and RBF were continuously recorded using PowerLab 8/35 (ADInstruments, CO) at a sampling rate of 1000 Hz for at least 20 minutes.

### Overnight urine collection

Mice were individually housed in metabolic cages overnight for urine collection, with free access to food and water. Urine volume was determined gravimetrically. Osmolality was measured using an osmometer (Wescor Vapor Pressure Osmometer VAPRO, model 5520), and sodium and potassium levels were assessed by flame photometry. Urine flow rate, and sodium and potassium excretion rates were normalized to body weight in grams.

### Acute volume loading

Under light isoflurane anesthesia, mice received subcutaneous injection of normal saline solution pre-warmed to 37°C, equivalent to 6% of their body weight. Urine excretion rate was assessed by placing the volume-loaded mice in metabolic cages, and urine samples were collected at 75-and 150-minutes post saline administration. This procedure was conducted both in the presence and absence of the selective vasopressin type 2 receptor (V_2_R) antagonist, tolvaptan (Tocris, Minneapolis, MN), administered at a dose of 1 mg/kg via intraperitoneal injection shortly before the subcutaneous saline loading.

### Plasma analysis

Mice were anesthetized and administered heparin before collecting blood from the left ventricle of the heart. After collection, blood was centrifuged at 4.5xg for 5 minutes at 4°C to separate plasma. Plasma osmolality was measured using an osmometer, while plasma copeptin was measured using a mouse copeptin ELISA kit by following the manufacturer’s instructions (LS BIO, Shirley, MA).

### Statistical Analysis

Data are presented as mean ± standard error of the mean (SEM). Sidak post hoc test was used following a two-way or three-way ANOVA for multiple comparisons between the experimental and control groups, where appropriate. All statistical analyses were performed using GraphPad Prism version 9 (GraphPad Software, San Diego, CA).

## Results

### Ovariectomy exacerbates blood pressure elevation in *Eln*^+/−^ female mice

Previously, we showed that *Eln*^+/−^ mice of both sexes exhibit higher nighttime systolic blood pressure (SBP) compared to their *Eln*^+/+^ counterparts^29^. We also reported that blood pressure elevation was relatively more pronounced in male *Eln*^+/−^ mice than in female *Eln*^+/−^, when compared to their respective controls, suggesting sexual dimorphism in the blood pressure phenotype resulting from *Eln* haploinsufficiency^29^. To test whether ovarian hormones play a role in the phenotypic differences, we examined diurnal blood pressure by radiotelemetry before and after OVX in *Eln*^+/+^ and *Eln*^+/−^ mice. Before OVX, *Eln*^+/−^ female mice exhibited higher nighttime SBP and slightly elevated diastolic blood pressure (DBP) compared to *Eln*^+/+^ mice, with no discernible difference in heart rate between the groups (Figure 1). Four weeks following OVX, nighttime SBP was slightly elevated in both genotypes, while daytime SBP was further elevated by ∼10 mmHg only in *Eln^+/−^* mice (Figure 1A and supplemental Figure 2A). In contrast, nighttime DBP showed a downward trend after OVX in *Eln^+/−^* mice, whereas daytime DBP was relatively unchanged in both genotypes (Figure 1B and supplemental Figure 2B). The augmented SBP elevation in OVX *Eln^+/−^*mice was confirmed to be independent of changes in locomotor activity, as both genotypes showed similar patterns in diurnal locomotor activity, before and after OVX (supplemental Figure 2C). Heart rate was similar between the two groups, with equal magnitude of decline following OVX (Figure 1C), while pulse pressure, an indicator of arterial stiffness, was elevated in both genotypes but appeared to be relatively more pronounced in *Eln*^+/−^ mice (Figure 1D). Together, these results indicate that ovarian hormones moderate systolic hypertension associated with *Eln* haploinsufficiency in female mice.

**Figure 1.**
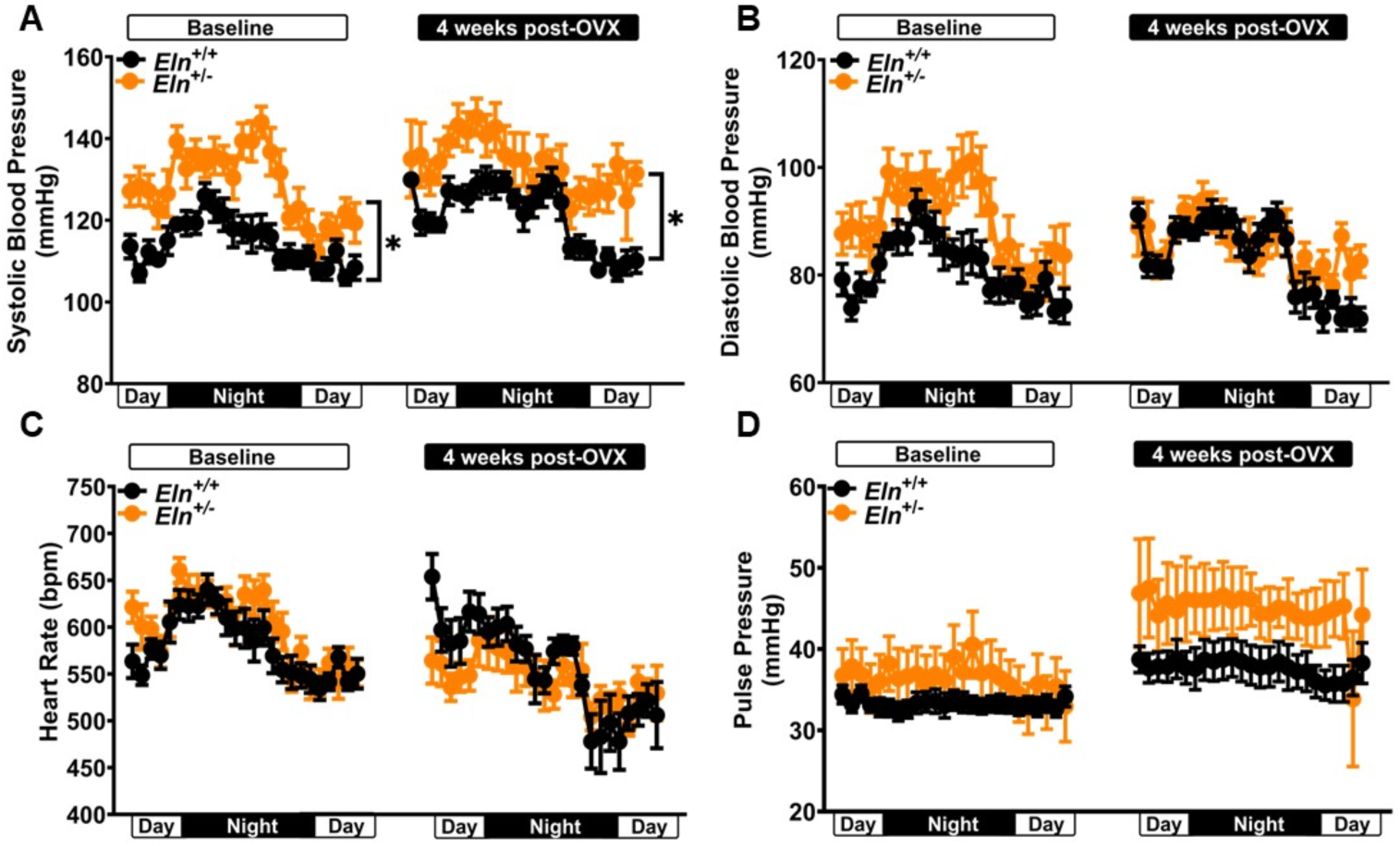
Diurnal blood pressure and heart rate of *Eln^+/+^* and *Eln^+/−^* mice before and after ovariectomy (OVX). (A – C) 24-hour systolic and diastolic blood pressure, and heart rate of *Eln^+/+^* and *Eln^+/−^*female mice before and 4 weeks after OVX. (C) A decrease in heart rate was observed in both groups after OVX. (D) Pulse pressure, an indicator of arterial stiffness, increased in both groups after OVX. Data are presented as mean ± SEM; Baseline *Eln^+/+^* (n=10), Baseline *Eln^+/−^* (n=13), OVX *Eln^+/+^* (n=8), OVX *Eln^+/−^* (n=8). \**P*<0.05, between-group comparison, analyzed using a two-way ANOVA mixed model with Sîdak post hoc multiple comparison tests.

### *Eln* haploinsufficient mice are insensitive to the anti-natriuretic effect of ovariectomy

Ovarian hormones, including estrogen and progesterone, have been shown to affect renal vascular tone and renal epithelial sodium transport through the activation of surface receptors and by their genomic effects^39–43^. The vascular and epithelial effects of ovarian hormones can alter renal perfusion and tubular function^44–46^. Therefore, we examined the effects of OVX on RBF and pressure-natriuresis relationship to determine whether *Eln* haploinsufficiency alters the effects of ovarian hormone depletion by OVX on renal hemodynamics and tubular function to exacerbate systolic hypertension. At baseline, sodium excretion rate trended high in sham *Eln*^+/−^ and OVX *Eln*^+/+^ mice, compared sham *Eln^+/+^* mice (Table 1, *P* = 0.098 and 0.063, respectively). However, both plasma and urine osmolality were similar between all groups. In sham mice, RBF trended slightly higher in *Eln*^+/−^ relative to *Eln*^+/+^ mice; However, RBF was substantially elevated in *Eln*^+/−^compared to *Eln*^+/+^ mice, following OVX (Figure 2A). Renal vascular resistance (RVR) was similar among the different groups but showed a decreasing trend in *Eln*^+/−^ mice after OVX (Figure 2B). GFR was similar between the two genotypes and trended downwards only in *Eln*^+/+^ mice after OVX, despite the augmented RBF in *Eln*^+/−^ mice after OVX (Figure 2C). In sham mice, filtration fraction (FF) trended lower in *Eln*^+/−^ mice compared to *Eln*^+/−^ mice and, following OVX, decreased further in both genotypes (Figure 2D). Furthermore, following OVX, the resulting daytime (Figure 2E) and nighttime (Figure 2F) pressure-natriuresis relationship showed a more rightward shift with little change in the slope in *Eln*^+/−^ mice compared to *Eln*^+/+^ mice. These trends were found to be consistent with increased sodium retention and salt-sensitive hypertension^13,47-51^.

**Figure 2.**
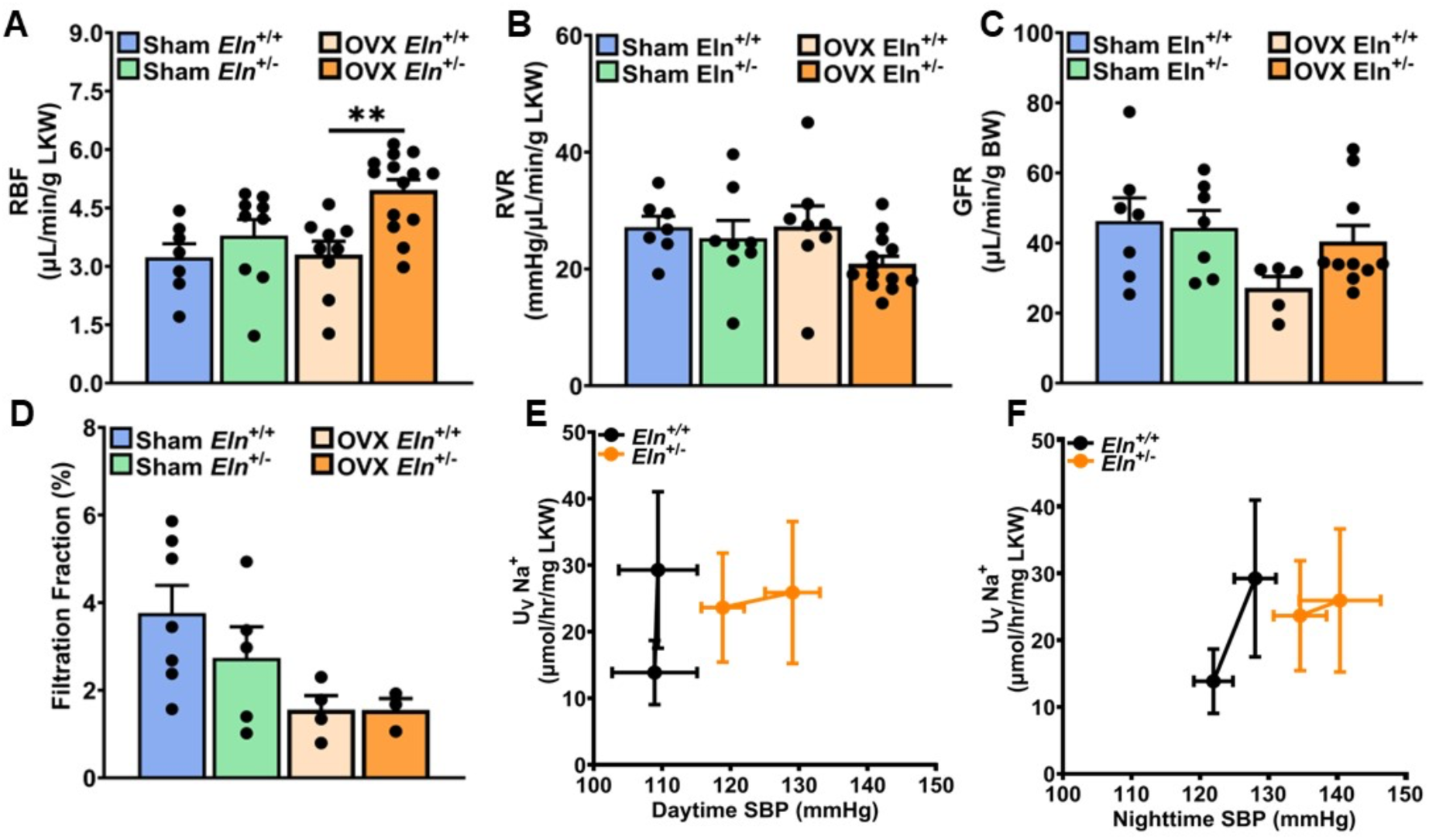
Renal hemodynamics and pressure-natriuresis in sham and ovariectomized (OVX) *Eln^+/+^* and *Eln^+/−^* mice. (A) Renal blood flow (RBF) was significantly increased in OVX *Eln**^+/−^*** mice, while (B) renal vascular resistance (RVR) was similar among the different groups but trended lower *Eln**^+/−^***mice after OVX. (C) GFR trended low in *Eln**^+/+^*** OVX mice but was similar among the other groups. (D) Filtration fraction (FF), calculated as the ratio of GFR to RPF, trended low in sham *Eln**^+/−^***mice and following OVX in both *Eln**^+/+^*** and *Eln**^+/−^***mice. Pressure-natriuresis curves generated with daytime (E) and nighttime (F) systolic blood pressure (SBP) and overnight urine sodium excretion rates of pre– and post-OVX in *Eln**^+/+^*** (n= 4-8) and *Eln**^+/−^***(n= 5–6) mice. Data are presented as mean ± SEM. UvNa***^+^***, urinary sodium excretion rate. LKW – Left kidney weight. \*\**P*<0.01

**Table 1:** Comparative analysis of urine flow rate, urinary Na^+^ and K^+^ excretion rates, and urine and plasma osmolality in sham and ovariectomized *Eln*^+/+^ and *Eln*^+/−^ mice. LKW, left kidney weight. Values are mean ± SEM.

|  | Sham |  | Ovariectomy |  |
| --- | --- | --- | --- | --- |
| Genotype | *Eln*<sup>+/+</sup> | *Eln*<sup>+/-</sup> | *Eln*<sup>+/+</sup> | *Eln*<sup>+/-</sup> |
| Number of animals | 6 | 7 | 6 | 9 |
| Urine flow rate (ul/hr/g BW) | 1.04 ± 0.20 | 1.35 ± 0.32 | 1.46 ± 0.33 | 1.70 ± 0.52 |
| Urinary Na<sup>+</sup> excretion rate (mmol/hr/g BW) | 182 ± 42 | 362 ± 85 | 386 ± 52 | 295 ± 74 |
| Urinary K<sup>+</sup> excretion rate (mmol/hr/g BW) | 695 ± 200 | 867 ± 240 | 807 ± 137 | 750 ± 235 |
| Urine osmolality (mmol/Kg) | 2833 ± 410 | 2263 ± 231 | 2709 ± 365 | 2151 ± 342 |
| Plasma osmolality (mmol/Kg) | 335 ± 20 | 340 ± 4 | 374 ± 27 | 374 ± 11 |

### Elevated systolic blood pressure in *Eln*^+/−^ mice is insensitive to low-salt diet

As previously reported, blood pressure elevation in *Eln*^+/−^ female mice is modest and insensitive to high-salt diet^29^. However, based on the observation herein that OVX enhanced SBP elevation in *Eln^+/−^* mice, and that overnight natriuresis in female *Eln^+/−^* mice phenocopied *Eln^+/+^* after OVX (Table 1), we postulated that blood pressure elevation resulting from *Eln* haploinsufficiency in female mice is due, at least partially, to abnormal sodium and water handling by the kidney. To test this hypothesis, we examined the effect of low-salt diet (LSD) on SBP in *Eln*^+/−^ female mice by placing sham and OVX mice on LSD for 10 days and reassessed diurnal blood pressure. As shown in Figure 3A, there was a slight downward trend in daytime SBP in sham *Eln*^+/+^ mice on LSD but no change in sham *Eln*^+/−^ cohort. In *Eln*^+/−^ OVX mice fed LSD, SBP remained elevated relative to their *Eln*^+/+^ cohort (Figure 3B). Conversely, DBP was similar in both sham and OVX groups of both genotypes on LSD (Figures 3C & 3D). Baseline heart rate was similar between the two genotypes, with or without OVX, and appeared to decrease at similar rates after LSD (supplemental Figure 3).

**Figure 3.**
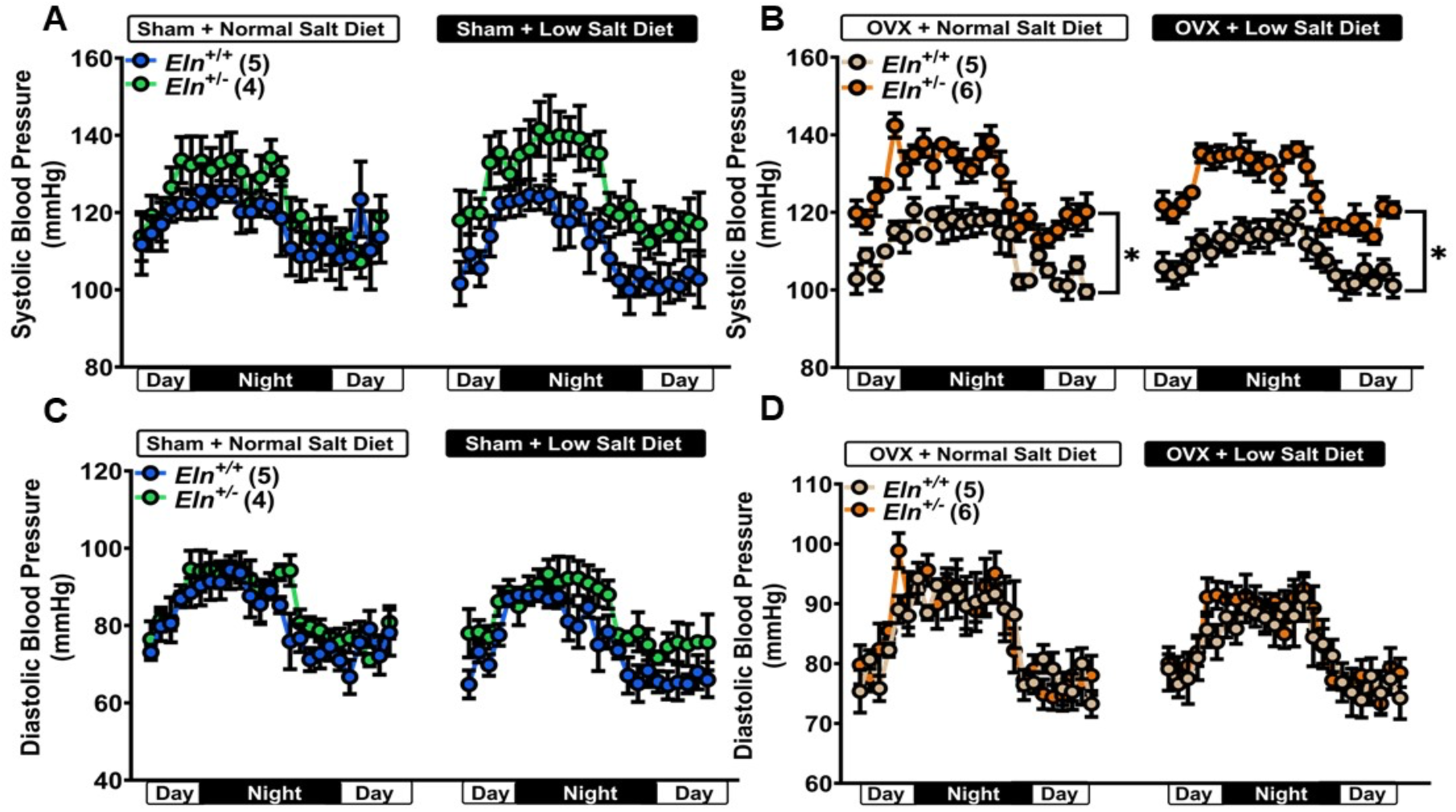
Effect of low-salt diet on diurnal blood pressure in sham and OVX *Eln^+/+^* and *Eln^+/−^*mice. Low-salt diet slightly lowered SBP in *Eln**^+/+^**,* thereby enhancing the difference in SBP between (A) sham and (B) OVX mice relative to their respective cohorts. Diastolic blood pressure in (C) sham and (D) OVX mice was similar. Data are presented as mean ± SEM, \**P*<0.05, between-group comparison, analyzed using a two-way ANOVA mixed model with Sîdak post hoc multiple comparison tests.

### Low-salt diet-induced augmentation of RAS activity is enhanced by OVX in *Eln*^+/−^ mice

Previous studies showed that RAS activity is augmented in *Eln*^+/−^ mice and contributes to the hypertensive phenotype at least partly by influencing vascular tone regulation^29,30^. However, whether augmented RAS activity by itself, or combined with the effects of ovarian hormones, enhances sodium reabsorption by the kidney, thereby contributing to sustained blood pressure elevation in *Eln*^+/−^ mice was unclear. Because LSD failed to lower SBP in OVX *Eln*^+/−^ female mice, and because LSD has been shown to induce a compensatory increase in renal sodium reabsorption via RAS activation^52^, we tested whether blood pressure insensitivity to changes in dietary sodium was due to maladaptive increase in RAS activity. We evaluated RAS activity by assessing blood pressure response to acute systemic blockade of angiotensin-converting enzyme (ACE) with captopril in conscious sham and OVX mice on LSD. In sham animals, SBP and DBP in *Eln*^+/−^ mice decreased by ∼30 mmHg within 30 minutes of captopril administration and remained low for more than 1 h post injection, whereas *Eln*^+/+^ mice exhibited a maximum blood pressure decline of ∼15 mmHg within 60 minutes of captopril injection (Figure 4A). The depressor response to captopril was accompanied by tachycardia in sham *Eln*^+/−^ but not *Eln*^+/+^ mice (Figure 4E). In OVX mice, LSD appeared to elicit a more robust and rapid depressor response to captopril injection in *Eln*^+/−^ (∼40-mmHg decline in SBP) relative to *Eln*^+/+^ mice (Figure 4B & 4D). As in sham animals, the depressor response to captopril was accompanied by tachycardia, which was less robust in *Eln*^+/−^ relative to *Eln*^+/+^ mice (Figure 4F). Overall, our data suggest that ovarian hormone depletion increases the sensitivity of the RAS to LSD and is further enhanced by *Eln* haploinsufficiency.

**Figure 4.**
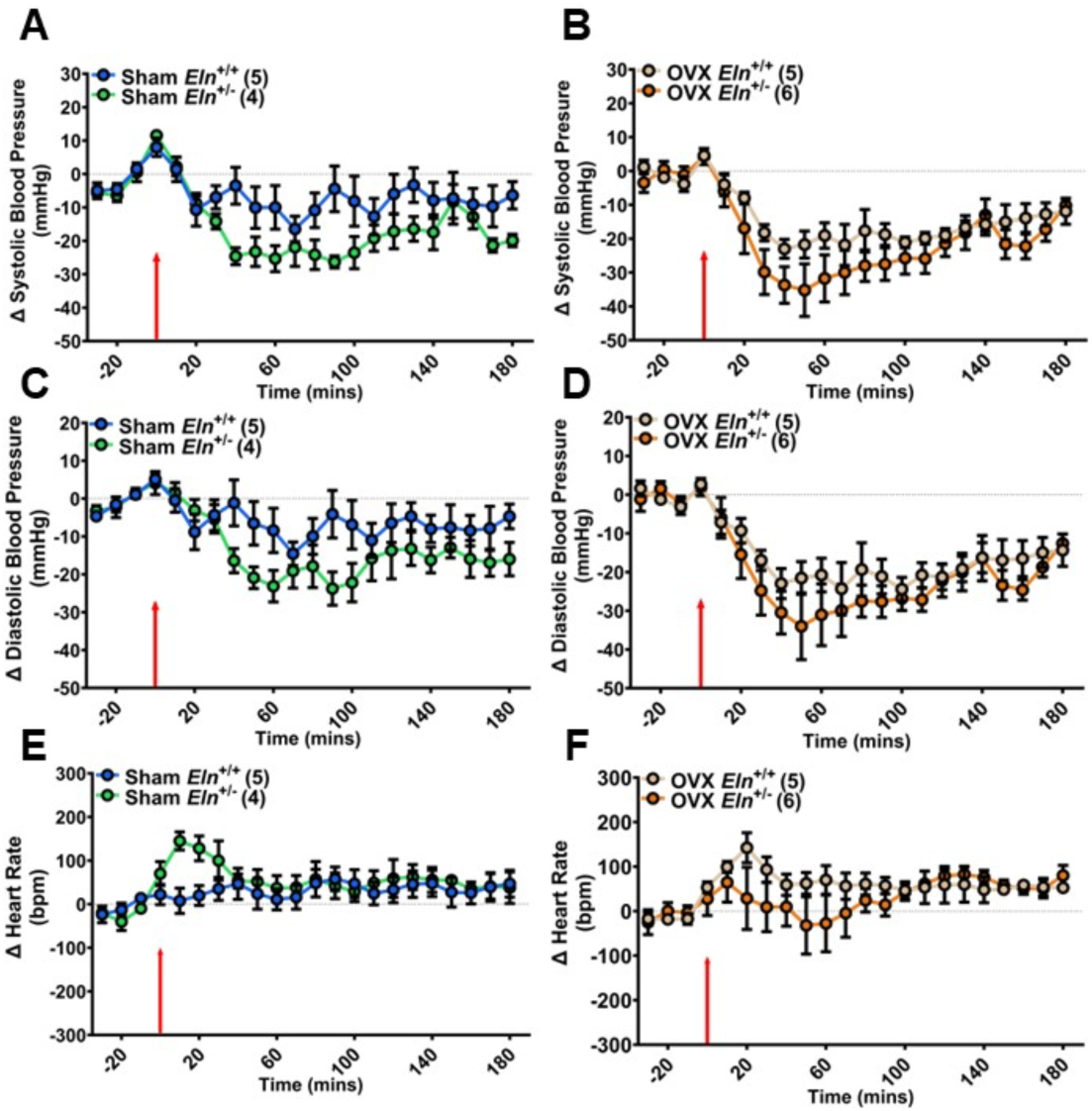
Blood pressure and heart rate response to acute systemic captopril administration in conscious mice instrumented with radiotelemetry devices. Baseline measurements were collected for ∼1 hour prior to captopril administration (10 mg/kg, i.p.). Average values over 30 minutes preceding the injection were subtracted from pre– and postinjection blood pressure and heart rate values. Δ Systolic blood pressure in sham (A) and OVX mice (B). Δ Diastolic blood pressure in sham (C) and OVX (D) mice. Δ Heart rate in sham (E) and OVX (F) mice. Data are presented as mean ± SEM. The time of injection is denoted by red arrow.

### Sustained elevation of SBP in OVX *Eln^+/−^* mice on LSD is associated with marked suppression of natriuresis and diuresis

To determine the basis of SBP elevation and blood pressure insensitivity to LSD in *Eln^+/−^* mice, we assessed urine flow rate and urinary sodium and potassium excretion rates in female mice on normal chow or LSD after sham and OVX. In sham *Eln*^+/−^ mice on normal salt diet (NSD), overnight urine flow rate was higher, while urine osmolality was lower compared to the respective sham *Eln*^+/+^ mice (Supplemental Figure 4A). Following LSD, urine flow rate increased markedly in the *Eln*^+/+^ mice, whereas it exhibited a decreasing trend *Eln*^+/−^ mice (Supplemental Figure 4A). Increased urine flow rate in *Eln*^+/+^ mice was accompanied by a decreasing trend in urine osmolality, whereas a lower urine flow rate in sham *Eln*^+/−^ mice was accompanied by a slightly increased trend in urine osmolality (Supplemental Figure 4C). In OVX mice, both urine flow rate and urine osmolality under NSD were similar between *Eln*^+/+^ and *Eln*^+/−^ mice. However, after LSD, urine flow rate showed a decreasing trend at similar rates in both genotypes, while urine osmolality appeared to trend differently (Supplemental Figure 4B and 4D). When compared to their sham counterparts on NSD, OVX increased urine flow rate with very little effect on urine osmolality in *Eln*^+/+^ OVX mice, and the effect of OVX on urine flow rate was slightly lowered by LSD (Supplemental Figure 4A vs 4B and 4C vs 4D). Conversely, both OVX and LSD did not appear to alter urine flow rate and urine osmolality under normal chow or LSD conditions in *Eln*^+/−^ mice (Supplemental Figure 4).

On NSD and prior to OVX, overnight sodium excretion rate in sham *Eln*^+/−^ mice showed an upward trend relative to values in sham *Eln*^+/+^ mice. However, following OVX, sodium excretion rate increased only in *Eln*^+/+^ relative to *Eln^+/−^*mice on NSD (Figure 5A). Conversely, LSD led to a 4-fold decrease in sodium excretion rate in sham *Eln*^+/+^ mice relative to ∼16-fold decrease in sodium excretion rate in sham *Eln*^+/−^ mice when compared to their respective NSD controls (Figure 5A). LSD equally lowered sodium excretion rate in *Eln*^+/+^ OVX mice compared to sham *Eln*^+/+^ control, whereas it slightly enhanced overnight sodium excretion rate in *Eln*^+/−^ OVX relative to *Eln*^+/−^ sham mice and to a similar level in *Eln*^+/+^ OVX mice (Figure 5A). *Eln* insufficiency or OVX had no effect on urinary potassium excretion rate in mice on NSD; However, LSD tended to lower potassium excretion rate in *Eln*^+/+^ OVX mice and both groups of *Eln*^+/−^ mice (Figure 5B). Plasma osmolality trended low after LSD in sham and OVX mice of both genotypes but were not significantly different from mice on NSD. Overall, these results suggest that i) LSD unmasks increased sodium reabsorption by the kidney due to *Eln* haploinsufficiency, and that ii) depletion of ovarian hormones minimizes the differences between *Eln^+/+^* and *Eln^+/−^*mice regarding renal sodium and water handling, implying that ovarian hormones mitigate renal tubular dysfunction resulting from *Eln* haploinsufficiency in female mice.

**Figure 5.**
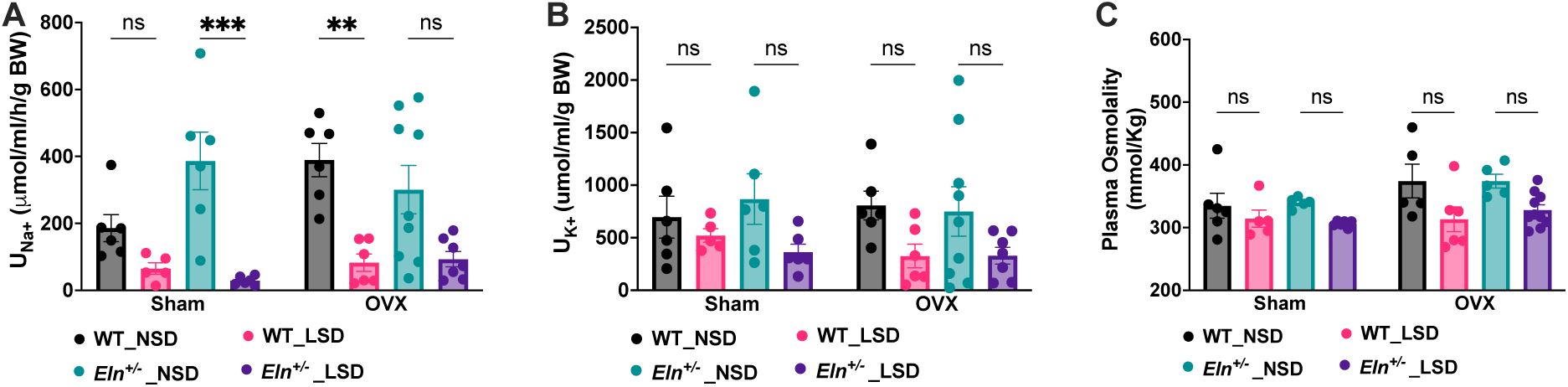
Low sodium diet exaggerates Na+ reabsorption in *Eln^+/−^* female mice. (A & B) The effect of low salt diet on overnight urine Na***^+^*** and K***^+^*** excretion rates in sham and ovariectomized (OVX) mice on standard chow (NSD) or low sodium diet (LSD). (C) The effect of LSD on osmolality of plasma obtained from sham and OVX mice at the end of 10 days on NSD or LSD. Data in bar graphs are expressed as mean ± SEM. Symbols represent individual animals in each experimental group and genotype. Between-group comparisons were analyzed using 3-way ANOVA mixed model with Sidak post hoc multiple comparison tests. **, \*\*\**P*<0.01, 0.001; ns, not significant.

### Amiloride reduces daytime SBP and promotes natriuresis in *Eln^+/−^* mice fed low salt diet

Elevated RAS activity increases blood pressure in part by promoting ENaC synthesis and insertion in principal cells of the distal nephron, thereby increasing sodium and water reabsorption^53^. As shown above, both OVX and *Eln* haploinsufficiency enhanced RAS activity in female mice. Therefore, to test whether sodium reabsorption via ENaC is involved in the exacerbation of hypertension upon ovarian hormone depletion in *Eln^+/−^* female mice, we examined the effect of subchronic administration of the ENaC inhibitor, amiloride, on diurnal blood pressure in OVX mice fed low salt diet. As shown in Figure 6, subchronic amiloride administration reduced SBP and DBP in both *Eln^+/+^*and Eln^+/−^ OVX mice. The effect of amiloride on blood pressure was more evident in daytime SBP of *Eln^+/−^* OVX mice, thereby narrowing the difference between the two genotypes in this period. Overnight urine sodium excretion rate trended higher in *Eln^+/−^* OVX mice following amiloride administration (Figure 6G). However, amiloride had no effect on urine osmolality, though it lowered urine potassium excretion rate equally in both genotypes after OVX (Figure 6H). These results suggest that increased sodium reabsorption via ENaC contributes to the exacerbation of blood pressure elevation in OVX *Eln^+/−^* female mice.

**Figure 6.**
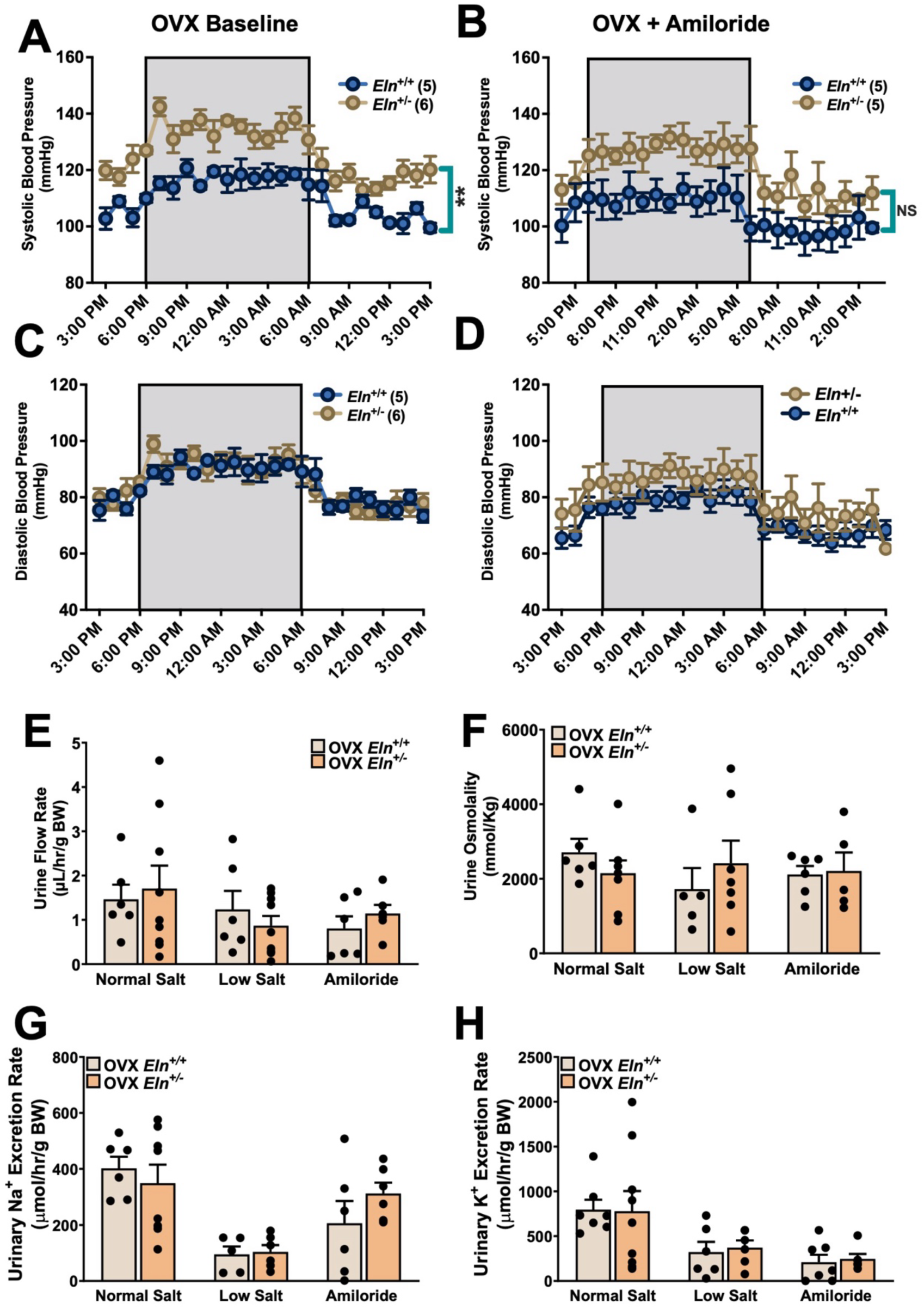
ENaC blockade with amiloride decreases SBP in OVX *Eln^+/−^* mice fed low salt diet. Amiloride was administered daily (1.5 mg/kg, i.p.) for 3 days at least 4 weeks after ovariectomy (OVX) and at the end of 10 days of low salt diet in wild type (*Eln**^+/+^***, n=5) and *Eln**^+/−^***(n=6) mice. Twenty-hour systolic (SBP, A & B) and diastolic (DBP, C & D) blood pressure was measured one day before and on the last day of amiloride injection. Overnight urine flow rate (E), osmolality (F), sodium excretion rate (G), and potassium excretion rate (H) before and after 3-day administration of amiloride in *Eln**^+/+^*** and *Eln**^+/−^*** OVX mice. Data points in bar graphs represent individual animals in each experimental group and genotype. Values in line graphs and bar graphs are expressed as mean ± SEM. \*\**P*<0.01, between-group comparison, analyzed using two-way ANOVA mixed model with Sîdak post hoc multiple comparison tests. NS, not significant.

### *Eln* haploinsufficiency suppresses excess sodium excretion in response to vasopressin receptor blockade after acute fluid volume expansion

The vasopressin system is widely recognized as the primary determinant of urine osmolality due to its regulation of water reabsorption in the collecting duct^54^. However, vasopressin can also affect sodium reabsorption by upregulating ENaC activity in the principal cells of the connecting tubule and collecting duct^55–57^. Indeed, certain mutations in vasopressin V_2_ receptor (V_2_R) that lead to increased ENaC activity are implicated in volume-dependent hypertension^56,58^. Therefore, we determined whether *Eln* haploinsufficiency promotes sodium retention by altering the effects of vasopressin on renal sodium and water handling. We examined urine flow rate and urinary sodium excretion rate following acute extracellular fluid volume expansion by a bolus administration of normal saline (6% of body weight, SQ), in the presence or absence of the V_2_R antagonist, tolvaptan (1 mg/kg, i.p.). In saline-treated sham and OVX *Eln*^+/+^ and *Eln*^+/−^ mice, urine flow rate was similar 75 minutes after acute volume loading (Figure 7A and 7B). Interestingly, fewer *Eln*^+/−^ mice produced urine at 75 minutes compared to their respective *Eln*^+/+^ controls (supplemental Figure 5). In the presence of tolvaptan, urine flow rate in sham and OVX *Eln*^+/+^ mice increased substantially at 75 minutes and was further elevated at 150 minutes. In contrast, urine flow rate in *Eln*^+/−^ mice was slightly elevated by tolvaptan only at 75 minutes in sham but not OVX animals (Figure 7A vs. 7B). In sham *Eln^+/+^* mice, tolvaptan increased urinary sodium excretion after acute volume loading, and this effect was abolished in *Eln^+/+^*OVX mice (Figure 7C). By contrast and similar to urine flow rate, tolvaptan had no effect on urinary sodium excretion rate in *Eln^+/−^*mice, after either sham or OVX (Figure 7C). These results indicate that *Eln* haploinsufficiency alters the sensitivity of the renal vasopressin system to acute sodium and water overload. The results also suggest that *Eln* haploinsufficiency enhances vasopressin signaling-mediated sodium, and thus water, reabsorption by the kidney.

**Figure 7.**
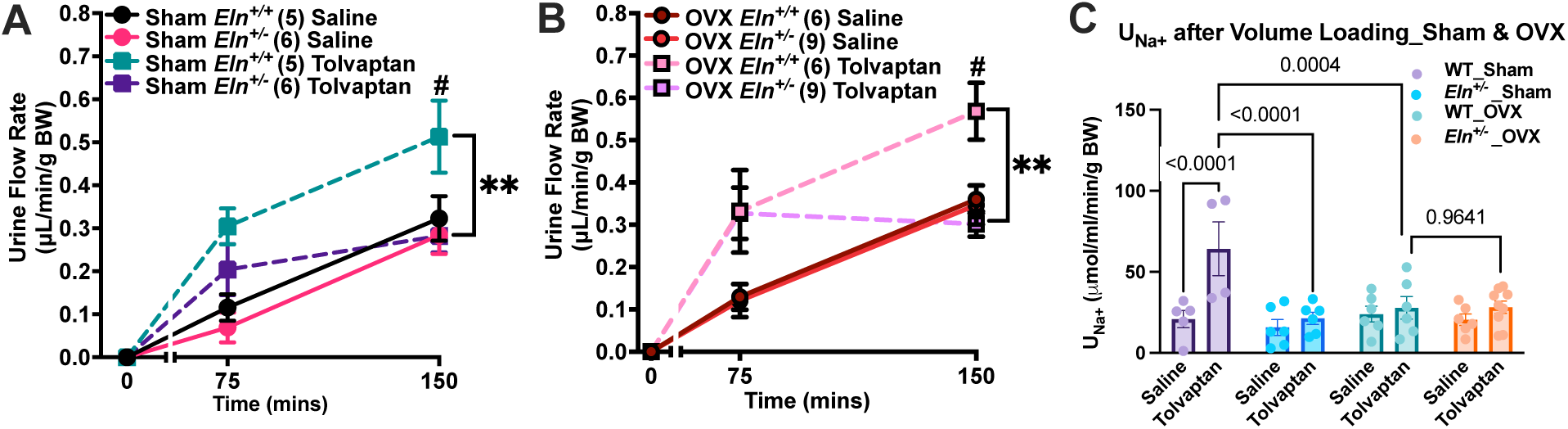
Urine flow and sodium excretion rate following acute extracellular fluid volume expansion with normal saline in the presence and absence of tolvaptan (1 mg/kg, i.p.). Urine flow rate after volume loading with saline, both with and without tolvaptan, are presented for sham-operated (A) and OVX (B) mice. (C) bar graph showing sodium excretion rate, which was calculated using the concentration of sodium in the urine collected from the 150th minute. Circles and solid lines represent urine flow rates without tolvaptan, while squares and dashed lines indicate rates with tolvaptan. Data are presented as mean ± SEM. \*\**P*<0.01, vs values at the 150th minute urine flow rates between *Eln*^+/+^ and *Eln*^+/−^ mice treated with tolvaptan. #*P*<0.05, urine flow rates between the 75th and 150th minute time points in mice given tolvaptan.

## Discussion

Elastin is expressed predominantly in the ECM of extensible organs such as the skin, lung, and conduit arteries, where it confers compliance^21,59^. In the kidney vasculature, *ELN* expression is confined to elastic fibers in small, multilayered preglomerular arteries^60^. Hemizygous deletion of *ELN leads to ELN* haploinsufficiency as part of microdeletion of several genes in a small region of human chromosome 7 causing Williams syndrome (WS), a rare genetic disorder with peculiar cardiovascular phenotype including hypertension and arteriopathy manifesting as arterial stiffening, elongation, and lumen narrowing^28,30,61-64^. Mice with heterozygous deletion of *Eln* phenocopy the cardiovascular pathology of *ELN* haploinsufficiency as occurs in WS, including elevated blood pressure and arteriopathy with remarkable vascular lamellar disorganization, thinning, and fragmentation in both conduit and resistance arteries^30,65,66^. Previously, we showed by radiotelemetry that blood pressure elevation in *Eln^+/−^* mice mimics isolated systolic hypertension that is more severe in male mice and liable to further elevation by high-salt diet^67^. Using this mouse model of *Eln* haploinsufficiency combined with in vivo methods and low sodium diet, we now show that ovarian hormones moderate blood pressure elevation resulting from *Eln* haploinsufficiency and that the pathophysiologic mechanism of sustained systolic blood pressure elevation in female *Eln^+/−^* mice, surprisingly, involves a change in renal vasopressin receptor sensitivity and ENaC activity, leading to enhanced sodium and water retention by the kidney.

The mechanisms by which the kidney maintains electrolyte and water balance are complex but key to long-term blood pressure control and are subject to regulation by several hormones, including ovarian hormones. Blood pressure modulation by ovarian hormones, mostly estrogen and progesterone, is mediated by the finetuning of tubular reabsorption of sodium and water^40,68^.

Previous studies investigating the effects of ovarian hormones on renal tubular sodium and water handling have produced conflicting results. On the one hand, studies show that, during periods of elevated circulating levels of estrogen, stimulation of estrogen receptors expressed in the distal nephron (connecting tubule and collecting duct) promotes sodium and water reabsorption by increasing the expression of apical AQP2 and ENaC and basolateral AQP1; these estrogen effects facilitate extracellular fluid conservation and increased cardiac output and blood pressure^11,40,69^. Estrogen receptor stimulation in these nephron segments in OVX rats has also been shown to upregulate ENaC expression, an effect that could be counteracted by progesterone treatment^70^. These effects of estrogen would favor blood pressure elevation by promoting sodium and water reabsorption. On the other hand, other studies in OVX rats reported that long term treatment with estrogen replacement suppressed adrenal Ang II receptor expression, thereby decreasing aldosterone release and ENaC expression, thus leading to reduced blood pressure^71^. The findings in the current study show that OVX elevates systolic blood pressure in both *Eln^+/+^* and *Eln^+/−^* mice, which is consistent with blood pressure lowering effect of ovarian hormones^72,73^. But more importantly, our study shows that, in the context of *Eln* haploinsufficiency, ovarian hormones dampen systolic blood pressure elevation by decreasing renal sodium and water reabsorption, at least partly, by modulating ENaC activity. Our finding is consistent with previous studies showing that estrogen has a downregulatory effect on RAS activity and ENaC expression in the kidney^40,74^. This effect of estrogen can lower blood pressure by decreasing sodium reabsorption in the distal nephron. Furthermore, the observed blood pressure lowering effect of ovarian hormones in female *Eln^+/−^* mice suggests a potentially additive or synergistic mechanism by which the effects of ovarian hormones and ELN (or its derivatives) in the distal nephron impinge on signaling pathways that mediate sodium reabsorption. Alternatively, the renal effects of ovarian hormones and ELN are plausibly redundant such that the blood pressure lowering effect of ovarian hormones might be a compensatory response triggered by *Eln* haploinsufficiency. In either mechanism, the renal effects of ELN appears to be protective or blood pressure lowering.

Furthermore, we made the surprising observation that *Eln* haploinsufficiency suppresses the excretion of excess sodium and water following acute ECFV expansion upon blockade of vasopressin V_2_R, even in the absence of ovarian hormones. As widely reported and well established, the urine concentrating effect of V_2_R activation in the collecting duct is mediated by the trafficking and insertion AQP2 that serves as water channel in the apical membrane of epithelial principal cells, with the hypertonicity of the medullary interstitium providing the driving force for water reabsorption^63,64^. Interestingly, our study suggests that *Eln* haploinsufficiency augments V_2_R-mediated water reabsorption in acute ECFV expansion. However, the most profound effect of OVX and *Eln* haploinsufficiency was on the retention of excess sodium that appears to be mediated by augmented V_2_R activity, which is a relatively less studied but widely recognized effects of V_2_R signaling on renal sodium reabsorption^55–57^. Several lines of evidence indicate that vasopressin can induce sodium reabsorption via ENaC in the distal nephron^55–58^. Accordingly, a previous study by Stockand and colleagues showed that chronic V_2_R activation promotes apical membrane ENaC insertion in collecting duct principal cells, thus enhancing sodium reabsorption and inducing Liddle syndrome-like hypertension^75^. The findings in the current study are consistent with these previous observations. Indeed, subchronic treatment of female *Eln^+/−^* mice with amiloride to block ENaC after low salt diet lowered systolic blood pressure, while slightly enhancing urine flow rate and urinary sodium excretion rate. These results are consistent with the hypothesis that chronic stimulation of vasopressin receptor signaling can cause blood pressure elevation by itself or work synergically with the RAS to promote sodium and water reabsorption, and thus volume-dependent hypertension.

This study sheds light on how ovarian hormones and altered sodium and water handling by the kidney are involved in hypertension associated with *Eln* haploinsufficiency. However, the specific mechanism by which ELN protein is physiologically involved in renal tubular function and, thus, regulation of water and electrolyte balance and long-term blood pressure control remain unexplored. It is conceivable that the modulatory effect of estrogen on vascular tone becomes more relevant in the context of elevated RAS activity as occurs in *Eln* insufficiency, such that depletion of ovarian hormones upon OVX further unmasks the elevated RAS activity. This is particularly plausible since *Eln* is not expressed in tubular epithelial cells (https://humphreyslab.com/SingleCell/displaycharts.php; https://esbl.nhlbi.nih.gov/KTEA/; https://cello.shinyapps.io/kidneycellexplorer/). Additionally, *Eln* haploinsufficiency might also enhance vasopressin signaling in the vasculature mediated by the low-affinity V_1_R to contribute to the augmented vascular tone and high blood pressure in *Eln* haploinsufficiency. These alternative mechanisms and hypotheses are not addressed by our study and remain to be explored.

## AUHTOR CONTRIBUTIONS

A.J.D, I.M., G.K., J.M., and P.O. performed experiments, and processed and analyzed data; A.J.D and P.O. drafted the manuscript; J.M. and P.O. edited and revised the manuscript; and P.O. approved the final version of the manuscript.

## ACKNOWLEDGEMENT

The authors thank Dr. Jeffrey R. Schelling for his constructive critiques of the manuscript.

